# Modification of gene expression after internalization of Growth Hormone (GH) into the cell nucleus

**DOI:** 10.1101/2023.02.07.525645

**Authors:** Gerard Morel

## Abstract

Growth hormone (GH) and many other hormones or growth factors have been shown to be rapidly internalized and translocated into the nucleus. The first event of peptide action is binding to its receptor which initiates both signal transduction pathways and internalization. The latter process involves the nucleus and probably gene transcription. In order to analyze the consequences of internalization of GH on gene expression, we used different populations of CHO cells, transfected with either full length GH receptor, or with defective receptor unable to trigger either signal transduction (deletion of box1) or the internalization of GH (Phe_346_ mutation in Ala). In addition, functional isolated nuclei were incubated 10 and 30 min with 50 nM GH in order to analyze the direct effect of GH on gene expression without surrounding cytoplasmic structures. The genes involved in signal transduction pathways were not revealed if GH internalization is the only functional activity in the whole cell as well as with isolated nuclei. In intact cell, internalization increased expression of 297 genes and decreased fewer than 10% of those known to be influenced by GH. Variations of expression in purified nuclei showed large variations with time. If cell signaling was not modified, cellular growth and proliferation, nucleic acid metabolism, cellular development, cell cycle and gene expression showed many variations with time. GH internalization shows direct effects on gene expression, different from those stimulated by signal transduction.

## INTRODUCTION

After the first description of internalization of [^3^H] labelled TRH to the nucleus (Gourdji et al, 1973) many others peptides have been detected in the nucleus of target cells (Rakowicz-Szulczynska et al, 1986; Smith and Jarett, 1987; Smith et al, 1988; Maher et al, 1989; Soler et al, 1989; Baldin et al, 1990; Jans, 1994; Morel, 1994; Wiedlocha et al, 1994, 1996; Maher, 1996; Jans et al, 1997) including Growth hormone (GH) and its receptor (GHR). After its binding to the plasma membrane receptor, GH is internalized in different cellular compartments (Roupas and Herington, 1989; Ilondo et al, 1992; Lobie et al, 1994). The internalization of the GH/GHR complex presumably involves clathrin-coated pits (van Kerkhof et al, 2001; Mertani et al, 2003) and/or caveolae (Lobie et al, 1994) and the ubiquitin conjugated complex. It has also been demonstrated that both GH (Lobie et al, 1991, 1994; Mertani et al, 1996) and GHR (Govers et al, 1997) are quickly internalized into the nucleus (Lobie et al, 1994). Indeed, [^125^I] labelled GH has been specifically found within pituitary nuclei and mitochondria *in vivo* as early as 30 min after GH injection (Mertani et al, 1996). Nuclear import requires recognition of the receptor-hormone complex by the cellular nuclear import machinery i.e members of NLS-recognizing IMP superfamily (IMPα/β heterodimer targets the internalized GH to the nucleus) (Jans et al, 2000).

In the same way, detection of GH in the mitochondria raised the question of its functional relevance in the regulation of oxidative phosphorylation as an essential cellular energy supplier. Consistently, it has been shown that GH can regulate the oxidative function of hypophysectomized rat liver mitochondria (Henneman, 1968; Isaksson et al, 1982; Davidson, 1987; D’Alessio et tal., 2005). More recently, we reported that GH can stimulate cellular respiration after binding to its receptor by initiating both a GH-dependent signal transduction pathway and the internalization of GH and GHR through a caveolar-dependent pathway (Perret-Vivancos et al, 2006). In these experiments, we demonstrated that the stimulatory effect was obtained through the GHR-signaling pathway involving the Box1 region, whereas inhibitory effects were reported after impairment of the signaling pathway. As the internalization pathway was functional, our results suggested a potential interaction between the signal transduction and the internalization pathways in regulating GH action on the cellular respiration. As we previously presented evidence for the presence of both GH and GHR within mitochondria (Ardail et al, 2010), the aim of the present study was therefore to discriminate between a possible direct effect of GH on the nucleus (a non-cell membrane GHR-mediated effect) and an indirect effect (a cell-membrane GHR-mediated effect) initiated at the plasma membrane through the activation of the GHR-dependent signal transduction pathway.

The main consequence of the simultaneous presence of GH/GHR in the nucleus is therefore expected to have an effect on gene transcription. In order to test this hypothesis, we used the transfected cell systems previously described (Perret-Vivancos et al, 2006) and isolated functional nuclei in order to analyse the consequences of GH/GHR internalization into nucleus in terms of genes expression using microarrays.

## MATERIALS AND METHODS

### Hormone and antibodies

Recombinant human GH (hGH) and a guinea pig anti-rGH antibody were obtained from Dr AF Parlow (National Hormone and Pituitary program, Harbour-UCLA Medical Center, Torrance, CA, USA). Reagents for cell culture were purchased from Invitrogen (Cergy Pontoise, France). The following antibodies were used in this work: a polyclonal IgG raised in guinea-pig against rGH; a monoclonal mouse antibody against GH receptor (mAb 263) that was a generous gift from Dr M.J. Waters (Queensland University, Brisbane); a goat IgG against guinea-pig IgG was obtained from DAKO (Copenhagen, Netherlands); and a goat IgG coupled to gold particles was from British Biocell International (Cardiff, UK).

### Cellular transfection of GH receptor cDNA

The expression plasmid, pIPB-1, containing the full-length cDNA of GHR (GHR_1-638_), under the transcriptional control of the human metallothionein IIa promoter and the simian virus (SV) 40 enhancer was constructed as previously described (Allevato et al, 1995). The construction of different GH receptor cDNA expression plasmids containing either a deletion of box 1 (GHR_Δ297-311_) or a single point mutation in position 346 (GHR_F346A_) has been previously described (Møldrup et al, 1991; Vanderkuur et al, 1994; Allevato et al, 1995). The cDNA constructions were transfected into CHO-K1 cells with lipofectin together with the pIPB-1 plasmid containing a neomycin resistance gene fused to the thymidine kinase promoter (Möller et al, 1992). Stable transfected cells were selected using 1 mg/ml G418 and transfectants were expressed into CHO cells that were further named CHO-GHR_1-638_, CHO-GHR_Δ297-311_ and CHO-GHR_F346A_.

### Hormone internalization

For hormone internalization kinetics, CHO cells were grown to ∼80% confluence in RPMI medium supplemented with 10% fetal calf serum, 50 units/ml penicillin and 50 µg/ml streptomycin before being serum deprived for 12 h. Cells were then incubated for 60 min with 50 nM rGH in serum free medium, then washed in the same medium, rinsed in PBS and subsequently fixed for 15 min in ice-cold 4% paraformaldehyde in PBS, pH 7.4. The cells were permeabilised in 0.1% Triton X-100 in PBS for 5 min, washed extensively with PBS and incubated for 1 h with a guinea pig anti-rat GH antibody diluted to 1/1000 (1 µg/ml) and the monoclonal mouse anti-rGHR (mAb 263) antibody diluted to 1/50 (25 µg/ml). GH and GHR detections were performed respectively with Cy3^™^ conjugated goat anti-guinea pig IgG diluted to 1/500 and Rhodol green^™^ conjugated goat anti-mouse IgG (Molecular Probes, Leiden, Netherlands) diluted to 1/100. Controls were performed by (a) omission of the primary antibody, (b) replacing primary antibody with the same protein concentration of preimmune mouse or guinea-pig serum and (c) preincubation (24 h at 4° C) of anti-rGH with 50 µg/ml of recombinant rGH or of mAb 263 with 20 µg/ml recombinant receptor extracellular domain.

### Confocal laser scanning microscopy (CLSM)

All images were acquired with a Zeiss LSM 410 confocal microscope equipped with laser sources, three separate detection channels with their own pinhole detector, and a fourth detection channel for DIC (Nomarski) transmitted light scanning microscopy. The fluorescence images of Cy3 and Rhodol green were collected sequentially on two different channels to avoid cross talk between fluorescence signals. Cy3 fluorescence was excited with a 543-nm HeNe laser source. A 560-nm dichroic beam splitter was used to separate laser excitation from emitted fluorescence, and an additional 590-nm long-pass filter was used after the pinhole in order to reject non-specific fluorescence. Rhodol green fluorescence was excited with a 488-nm Argon laser source. A 510-nm dichroic beam splitter was used to separate laser excitation from emitted fluorescence and an additional 515-545 nm band-pass filter was used after the pinhole to reject non-specific fluorescence. The pinhole aperture was set to one Airy disk unit for each channel. An image of the same field was recorded in DIC transmitted light in order to facilitate the cell and nuclear contour delineation in the further image analysis stage. Image averaging (Kalman filtering) was used for the acquisition of the fluorescence images to improve the signal-versus-noise ratio.

### Analysis of CLSM images

Ten image triplets (Cy3 – Rhodol green – DIC) were recorded per experimental slide. The images were stored on a computer disk and analyzed with a SAMBA 2005 image analysis system (SAMBA Technologies, Meylan, France). Dedicated software was developed for this particular application. Briefly, the DIC image of a triplet was used to interactively extract the contours of the cells and of the cell nuclei. From these contours, the program built the cytoplasm and nucleus binary masks of each cell. Then the Cy3 image was loaded. For each cell, the fluorescence intensities (arbitrary unit) were measured using the previously obtained binary masks inside the cytoplasm and the nucleus. Two parameters were obtained: the integrated fluorescence, which is the sum of intensity values measured within the mask; and the average fluorescence intensity, which is the integrated fluorescence divided by the area (in pixels) of the binary mask. Next, the Rhodol green image was loaded and the same measurements were made using the binary masks as previously described (Mertani et al, 2003).

### Preparation of functional nuclei

CHO-GHR_1-638_ cells were rinsed in PBS as previously described. Nuclei were prepared by using nuclei pure prep nuclei isolation kit (Sigma Aldrich Chemicals, Saint Quentin Fallavier, France). Nuclear fractions were incubated for 10, 30 min with 50 nM of rGH. Immuno-electron microscopy and microarrays were performed as described below.

### Immuno-electron microscopy

Purified nuclei were prepared as described above. The nuclear suspension was spun at 8 500 *g*. The pellet was washed in 0.1 M phosphate buffer, pH 7.4, at 4° C and fixed in 4% paraformaldehyde, 0.1% glutaraldehyde in 0.1 M phosphate buffer, pH 7.4. The samples were embedded in LR-White resin (Sigma Aldrich). Polymerization was performed in gelatine capsules at 60° C for 24 h.

Ultrathin sections of 70 nm were obtained using an Ultracut S (Leica, Vienna, Austria). Immunocytological detection of GH was carried out using a guinea pig anti-GH IgG (dilution 1/100 in 100 mM phosphate buffer, 300 mM NaCl, 0.05% Tween-20, 0.5% BSA, pH 7.4) and revealed by an anti-guinea pig IgG conjugated with 10 nm gold particles (British BioCell, Cardiff, U.K) at 1/50 in 20 mM Tris, 300 mM NaCl, 0.05% Tween-20, 0.5% BSA, pH 8.2. The contrast was performed by incubation in 4% uranyl acetate pH 7 for 30 min. Electron micrographs were taken with a Philips CM120 electron microscope, operating at 80 kV (Centre Technologique des Microstructures, Villeurbanne, France).

### Microarrays

#### RNA extraction

Total RNA from three independent experiments of the cells or nuclei were prepared using the RNeasy mini-kit (Qiagen) according to the protocol provided by the manufacturer, including the additional step of DNAse treatment. Total RNA yield was measured using OD_260_ (optical density), and the quality of isolated total RNA was evaluated with the Agilent 2100 bioanalyzer (Agilent Technologies, Palo Alto, CA, USA) for microarray analysis or on a 1% ethidium bromide stained-gel for quantitative PCR.

### Microarray analysis

#### RNA amplification

Total RNA, 2 µg, was amplified and labelled by a round of *in vitro* transcription using the message amp aRNA kit (Ambion, TX, USA) following the manufacturer’s protocol. Before amplification, all tubes were spiked with synthetic mRNA at different concentrations in order to verify the quality of the amplification. aRNA yield was measured with an U.V. spectrophotometer and quality verified on nanochips with the Agilent 2100 bioanalyzer (Agilent Technologies).

#### Array hybridization and processing

Biotin labelled aRNA (10 µg), were fragmented with 5 µl fragmentation buffer in a final volume of 20 µl. Fragmented aRNA were added to the hybridization solution (GE Healthcare Europe GmbH, Freiburg, Germany) in a final volume of 260 µl and injected into the CodeLink Uniset Mouse I Bioarrays containing 10,000 mouse genes probes (GE Healthcare). Arrays were hybridized overnight at 37° C at 300 rpm (8 *g*). The slides were washed in a stringent buffer containing 0.1 M Tris-HCl pH 7.6, 0.15 M NaCl and 0.05% Tween 20 (TNT) at 46 °C for 1 h followed by a streptavidin-Cy5 (GE Healthcare) detection step. Each slide was incubated in 3.4 ml streptavidin-Cy5 solution for 30 min. Thereafter, the slides were washed four times in 240 ml TNT buffer. For the final washes, the slides were rinsed twice in 240 ml water containing 0.2% Triton X100. Slides were finally dried by centrifugation at 600 rpm (32 *g*).

Slides were scanned using a Genepix 4000B scanner (Axon Instruments, Union City, CA, USA) and Genepix software, with the laser set at 635 mm, laser power at 100% and photomultiplier tube voltage at 60%. The scanned image files were analyzed using CodeLink expression software, version 4.0 (GE Healthcare), which produces both raw and normalized hybridization signals for each spot on the arrays.

#### Microarray data analysis

The overall raw hybridization signal intensity of arrays was normalized using CodeLink expression software version 4.0 (GE Healthcare) by setting the raw hybridization signal on each array as ratio to the median of the array (median intensity is 1 after normalization) for better cross-array hybridization. This study used normalized signal intensities. Probes containing incomplete data were eliminated from the list. The threshold of detection was calculated using the normalized signal intensity of the 100 negative controls represented in the array. Spots with signal intensity below the threshold were termed ‘absent’.

Quality of processing was evaluated by generating scatter plot representations of positive signal distribution. Signal intensities were then transformed by logarithm (base 2). A differential expression of at least 1.3 (minimal fold change) was used as a threshold for generating the final list of genes modified by GH, whereas the average fold change is used when describing individually the expression level of genes of interest. The minimal fold change is indicated in results. Statistical comparison and filtering were achieved by using the Genespring software 7.0 (Agilent Technologies).

## RESULTS

### GH-GHR localization in cells

In order to determine the consequences of the nuclear internalization of GH on gene transcription, we have used different cell populations transfected with different GHR constructs. Figure 1 illustrates the GH internalization process 30 min after addition of GH in CHO cells stably transfected with the complete rat GHR cDNA (CHO-GHR_1-638)_. Although similar results were observed (Figure 1B) after deletion of Box1 (CHO-GHR_Δ297-311_), the mutation of phenylalanine 346 residue in alanine (CHO-GHR_F346A_) resulted in a serious alteration of GH internalization (Figure 1C).

**Figure 1:**
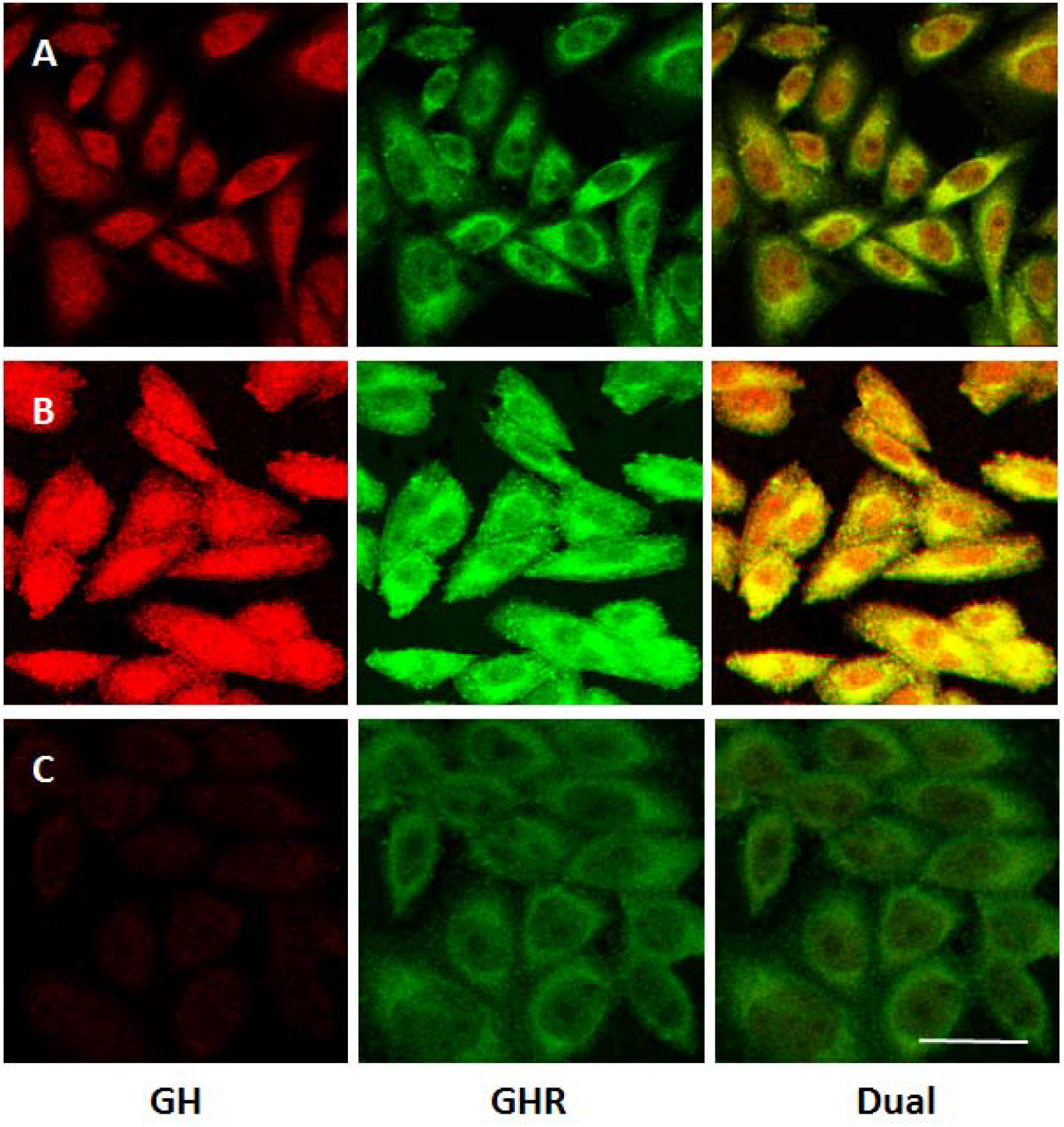
GH and GHR internalization and nuclear translocation in CHO cells expressing a GHR construct (A) the full length GHR (CHO-GHR_1-638_); (B) the GHR deleted of box 1 (CHO-GHR_Δbox1_); (C) with a point mutation of phenylalanine 346 (CHO-GHR_F346A_). CSLM analysis was performed at 30 min after 50 nM rGH stimulation as described in <u>Materials and methods</u>. The localization of GH is exposed in the left hand column, the GHR in the middle column and the dual localization (surimposed images) in the right hand column. Internalization process of GH disappeared after mutation of Phe 346. Scale bar: 10 µm.

### Effects of GH on gene expression

The consequence of GH incubation revealed a total of 125 increased and 3 decreased genes with a greater than 1.3 fold change in Box 1 deleted transfected cells (CHO-GHR_Δ297-311_) compared to controls (CHO-GHR_1-638_), and 279 overexpressed and 28 repressed genes in mutated transfected cells (CHO-GHR_F346A_) compared to controls. These genes were classified according to their molecular function.

Genes whose transcription in response to GH-dependent signal transduction pathways (CHO-GHR_1-638_ *vs* CHO-GHR_Δ297-311_) could be sorted mainly in 8 (Table 1) over 25 categories. Among them, one categorie confirms signal transduction of GH. Twelve genes were directly activated by GH after binding to its receptor on the plasma membrane: in particular, genes involved in GH signaling, ERK/MAPK and IGF-1 signaling (Table 1). Others genes activated by GH are involved in cell death [12], cellular assembly [14], cellular growth and proliferation [10], cellular development [9], protein synthesis [4] and the biochemistry of small molecules [9]. Collectively these results underline the involvement of the GH-dependent signaling pathways in both cell function and maintenance.

**Table 1:** List of genes modulated by GH membrane signal transduction in CHO cell

| Function | Number of genes regulated (up/down) | Symbol |
| --- | --- | --- |
| Cellular Growth and Proliferation | 34/0 | SH2B3, HDAC10, PBK, JUN, CSPG4, POSTN, LAMB1, AKT3, SLC9A3R2, SERPINE1, SLC12A4, DTL, ENPP1, RRAD, DCN, FXYD1, STAT3, MAFF, DES, CDK1, HDAC5, IVNS1ABP, GPR56, IMMT, B3GNT3, IRS1, CXCL12, MAZ, MAPK10, ZYX, UBTF, SFN. |
| Gene Expression | 26/0 | SH2B3, SNAPC3, HDAC10, ARNT2, VRK1, TCEB2, JUN, PMF1, SMARCC2, ATN1, HOXA2, DCN, FXYD1, HIVEP3, STAT3, MAFF, CDK1, HDAC5, IVNS1ABP, GMEB2, SREBF2, FOXN3, MAZ, RPS6KA4, UBTF, BCL9L |
| Cellular Development | 24/0 | SP6, SH2B3, RRAD, DCN, RHOJ, STAT3, MAFF, HDAC5, IVNS1ABP, JUN, CSPG4, IRS1, CXCL12, MAPK10, POSTN, LAMA3, AKT3, ZYX, UBTF, SFN, DOCK3, SERPINE1, MCPT4, DMBT1. |
| Cell-To-Cell Signaling and Interaction | 16/0 | SH2B3, BOC, DCN, PRSS3, STAT3, PPFIBP1, RS1, GPR56, CXCL12, IRS1, LAMB1, LAMA3, POSTN, ZYX, DOCK3, SERPINE1. |
| Small Molecule Biochemistry | 14/0 | ADSSL1, ENPP1, UGT1A6, ACOX1, MS4A1, STAT3, DES, RDH16, ADSS, IMMT, CXCL12, MAPK10, MSRA, GLDC. |
| Cellular Assembly and Organization | 14/0 | DES, CDK1, MCM4, HDAC5, JUN, IRS1, CXCL12, LAMA3, MAPK10, ZYX, SLC9A3R2, SMARCC2, SFN, UBTF. |
| Molecular Transport | 8/0 | RDH16, JUN, ACOX1, CXCL12, FXYD1, AKT3, UBTF, SFN |
| Cell Death | 19/0 | BRD2, DCN, STAT3, PBK, PSIP1, CDK1, PROKR1, HDAC5, AQP3, JUN, CLK3, SREBF2, CXCL12, IRS1, LAMA3, MAPK10, AKT3, SERPINE1, SFN. |

The main classes of genes involved after GH/GHR nuclear internalization (CHO-GHR_1-638_ *vs* CHO-GHR_F346A_) are reported in Table 2. No change in the expression of genes involved in signal transduction was noticed with this GHR construct. Through the clusters, the sole gene mostly stimulated was GHR mainly after GH internalization.

**Table 2:**
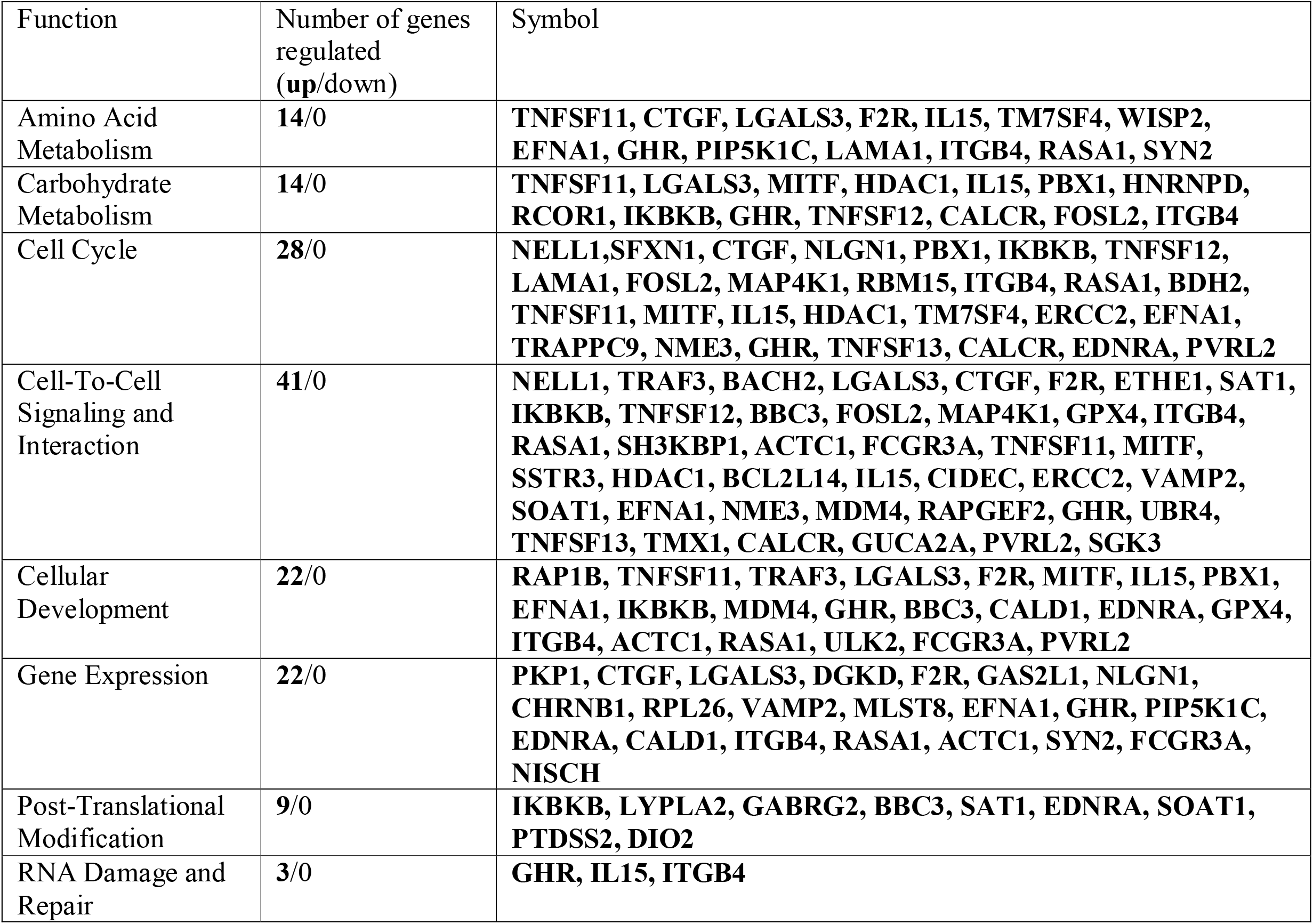
List of genes modulated by GH translocation to nucleus in CHO cells

After internalization, 44 genes are activated in the cluster of cell death. Then, the most important gene clusters activated by GH internalization are involved in cell cycle [20], cellular growth and proliferation [12]. Collectively, our results strongly underline that both nuclear GH internalization and the GH-dependent activation of signal transduction pathways are able to modulate different gene expression clusters.

### *GH localization in* isolated nuclei

At the electron microscopic level, GH can be detected within nucleus 30 min after incubation of isolated nuclei with the hormone (Figure 2). Without prior incubation with GH, GH immunoreactivity was still able to be detected in the nuclear fraction (Figure 2A), but the intensity of the signal (gold particle) strongly increased in purified nuclei after GH addition (Figure 2B).

**Figure 2:**
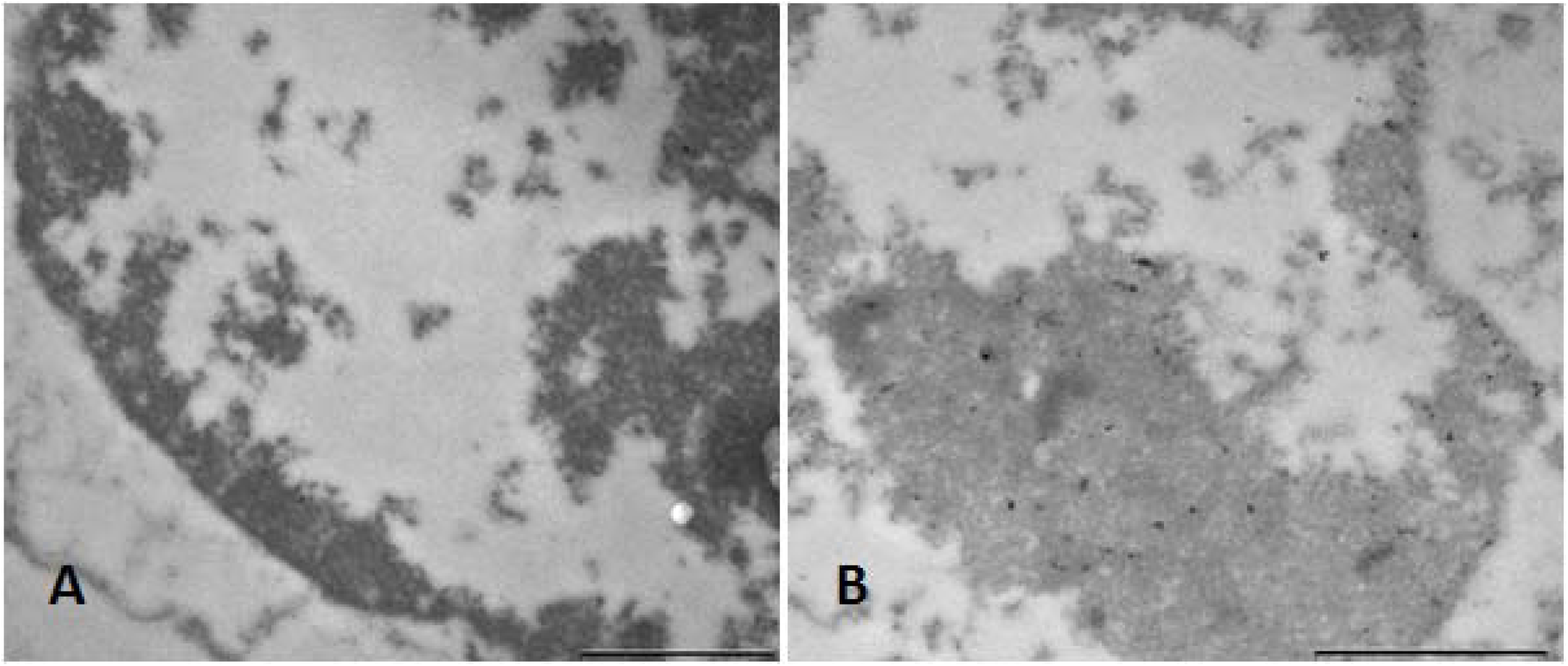
Nuclear internalization of GH into functional isolated nuclei. A: control nucleus: some gold particles are visualized on chromatin, B: after 30 min of GH incubation numerous gold particles are observed. Scale bar: 50 nm.

### Effect of GH on gene expression in isolated nuclei

The clusters of genes that have been modulated (fold change 1.3), are similar regardless of the incubation time (10 and 30 min), with exception for cluster containing only 1 gene. In this way, data observed were similar in comparison with clusters revealed by analysis of GH internalization in cells.

The time course experiments on the effect of GH on gene transcription in isolated nuclei are summarized in Tables 3 and 4. When functional isolated nuclei were incubated for 10 min in the presence of 50 nM GH, 54% of genes expression modified by GH addition were found to be up-regulated whereas 42% down-regulated (Table 3). For a longer incubation time (30 min) with GH, changes in genes expression profiles were noticed: only 8% of genes that increased during the 10 first min showed similar variation whereas 27% were not modified and 85% were decreased. By contrast, among the genes whose expression decreased during the 10 first min after GH exposure, 10% showed similar variation after 30 min of incubation with the hormone, 57% did not show any variation whereas 33% increased. Among the genes whose expression were unchanged during the 10 first min of incubation with GH, 18% increased, 31% decreased and 51% were not modified 30 min after GH exposure.

**Table 3:**
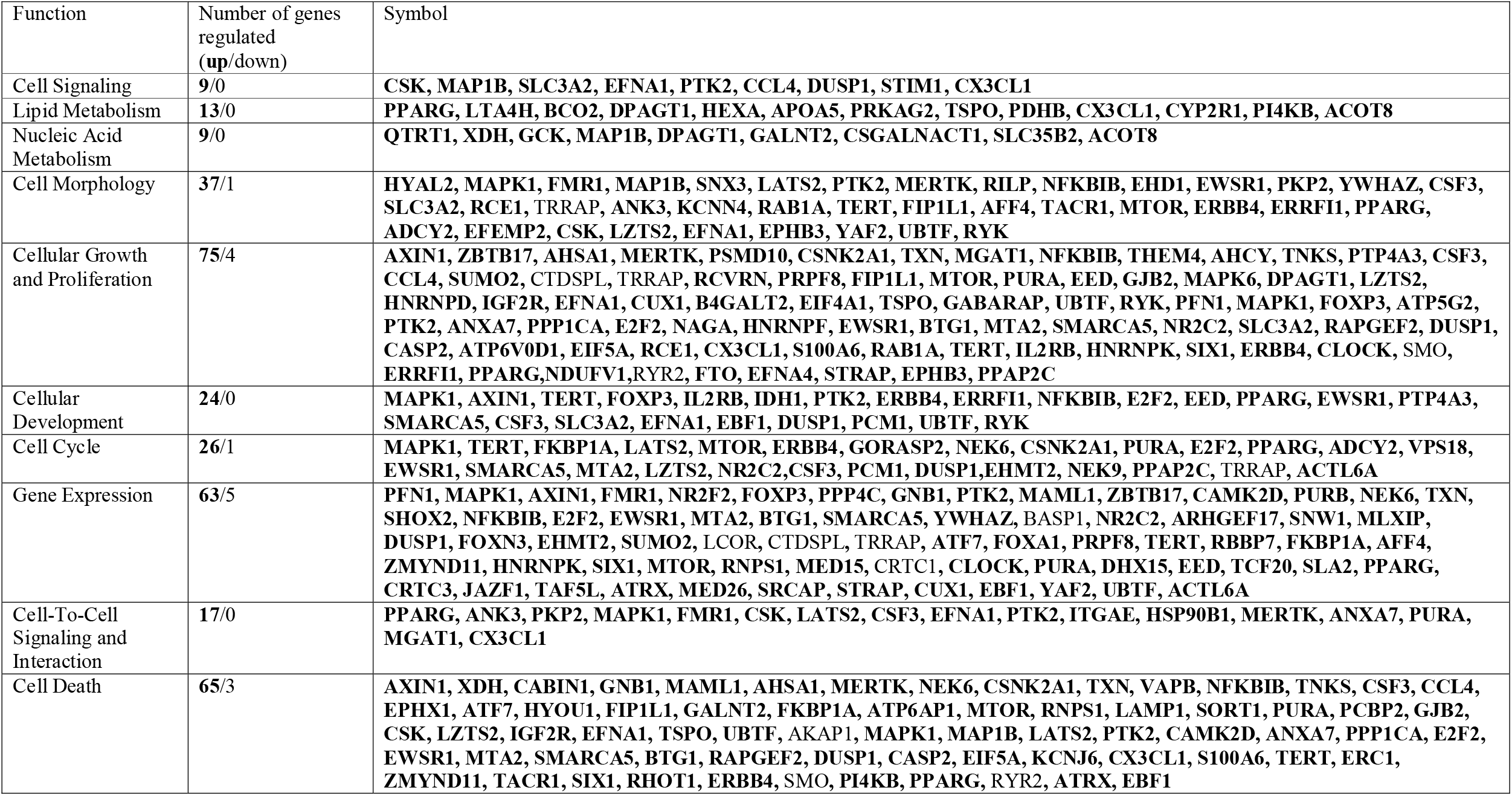
List of genes modulated by 10 min GH incubation with isolated nucleus

**Table 4:**
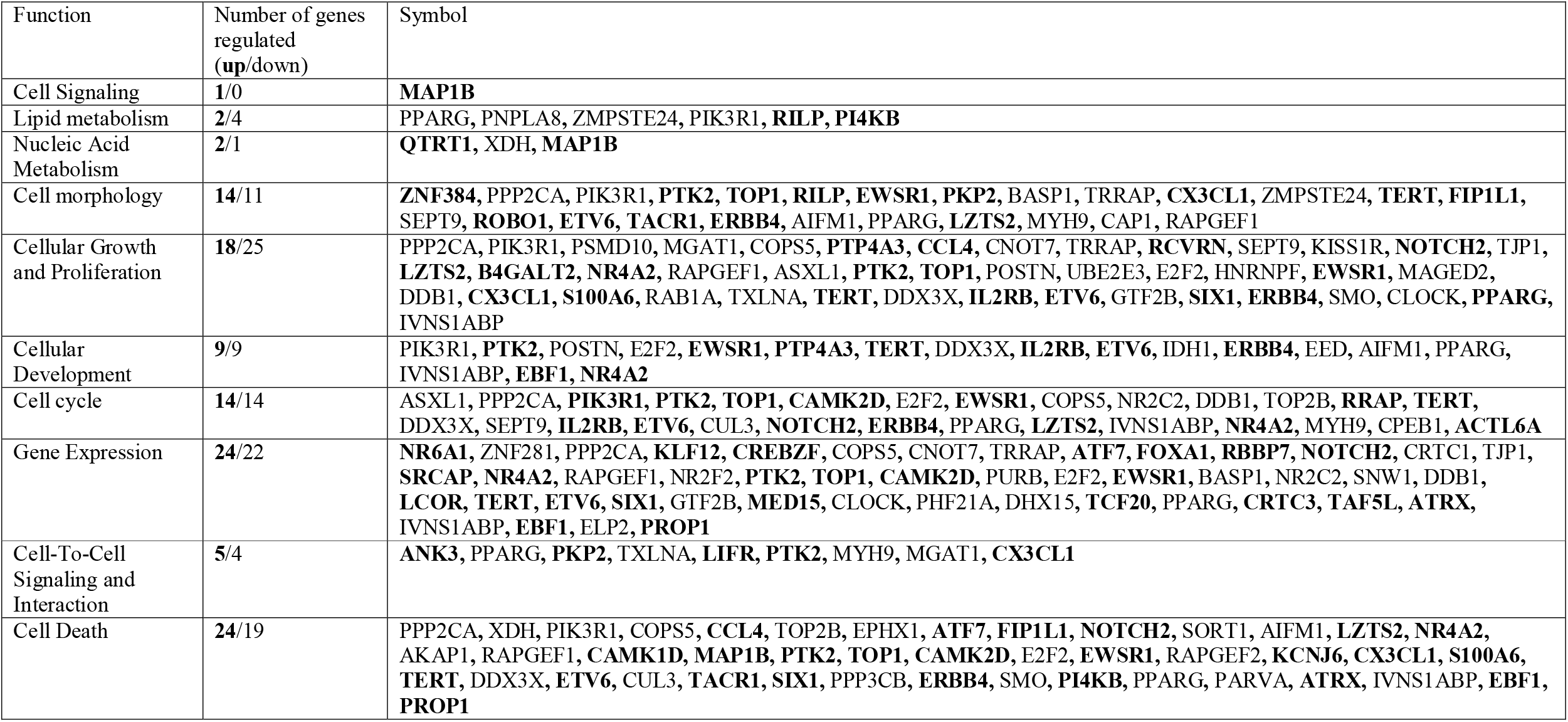
List of genes modulated by 30 min GH incubation with isolated nucleus

Collectively, these results demonstrate firstly that GH has a direct effect on isolated nuclear fractions and secondly that this effect on panels of different genes depends on the time of exposure to GH (10 min in Table 3 and 30 min in Table 4). The differential effects of GH relate to basic cellular functions such as cell signaling; lipid metabolism, nucleic acid metabolism; cell morphology; cellular growth and proliferation; cellular development; cell cycle; gene expression, cell-to-cell signaling and interaction and cell death, respectively.

### Overview of genes simultaneously modified in CHO cells and isolated nuclear fractions

As reported in Table 5, only 15 genes were found to be simultaneously modified in isolated nuclei and in whole CHO cells after incubation with GH. These genes are mainly involved in cell to cell signaling and internalization (BOC, TNFSF12, MAP4K1, CSK) or cell cycle (TNFSF12, MAP4K1).

**Table 5:**
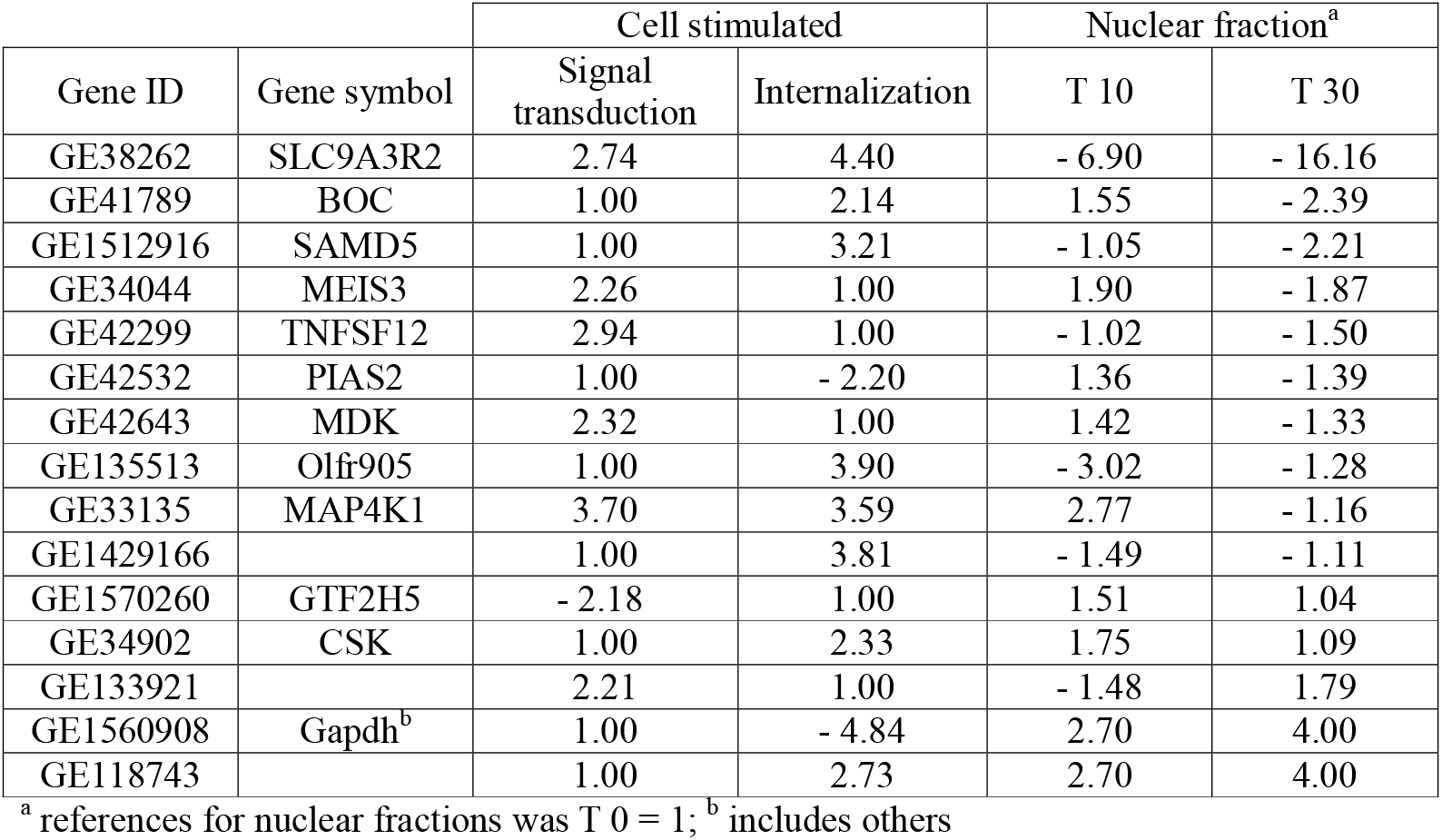
List of gene modulated by GH in cell or nuclear fraction simultaneously

## DISCUSSION

As with many other hormones such as insulin, fibroblast growth factor or prolactin (Hawker and Granger, 1994; Jans, 1994; Shah et al, 1995; Jans and Hassan, 1998; Johnson et al, 1998; Jans et al, 2003), GH can be internalized into the nucleus of target cells. The conclusion of these studies of internalization processes was limited in terms of potential effects on gene transcription. For GH, recent data have shown that nuclear GH internalization in a regenerative liver model was strongly correlated with the proliferative status of cells (Conway-Campbellet al, 2007). Other microarray analysis of the *in vivo* effect of hypophysectomy and GH treatment reveal transcript changes resulting from severe hormonal deficiency (hypophysectomy) and chronic treatment of these hypophysectomized animals with human GH also caused significant changes in gene expression patterns (Flores-Morales et al, 2001). However, it was very difficult in this study to correlate the variations in the expression of genes to specific physiological change since the variations observed resulted from physiological changes due to GH in absence of other pituitary hormones.

In order to determine the role of GH after its internalization in the nucleus of CHO cells, a differential study of gene expression profiles was carried out with two cellular transfection models displaying a modification of the internalization process of GH (after mutation of phenylalanine/alanine 346) (Allevato et al, 1995), or a modification of the specific signal transduction pathway (after deletion of box1) (Vanderkuur et al, 1994), respectively. These models of cells have been previously used to study the kinetic of GH internalization (Ardail et al, 2010). These three models of cells show positive GH nuclear internalization in CHO-GHR_1-638_ and CHO-GHR_Δ297-311_, but not in CHO-GHR_F346A_. Modification of signal transduction was observed only in CHO-GHR_Δ297-311_ (Ardail et al, 2010). Internalization and signal transduction are independent events regarding their initiation. Cells expressing the GHR after deletion of the Box1 region were unable to modify the expression of any gene involved in signal transduction pathways. Inversely, cells expressing the GHR mutated on the Phe 346 residue did not internalize GH whereas modifications of the expression of genes involved in signal transduction were observed. Given our experimental approach, we are able to differentiate GH effects due to signal transduction or internalization process, respectively. The impact of signal transduction pathways after exposure of cells to GH was studied by differential gene expression using CHO-GHR_1-638_ *vs* CHO-GHR_Δ297-311_, which showed activation of the pathway JAK/STAT5, characteristic of GH signal transduction (Argetsinger and Carter-Su 1996; Moutoussamy et al, 1998). In contrast, no modification of signal transduction from differential expression CHO-GHR_1-638_ *vs* CHO-GHR_F346A_ was detected, suggesting that internalization of GH do not alter signal transduction. Additionally, the results obtained with isolated nuclei incubated with GH show that the genes involved in the regulation of the signal transduction pathways initiated at the plasma membrane were not modified whatever the time of contact with the hormone. The direct action of GH in isolated nuclei shows similar consequences as functional GH internalization without signal transduction (CHO-GHR_Δ297-311_).

Many integral membrane proteins have been previously reported to be translocated to the nucleus through many hypothetical mechanisms and the biological consequences of this translocation are currently under investigation. Regarding the nuclear trafficking of EGFR family proteins, endocytosis and endosomal sorting are both required for the nuclear transport of the EGFR complex (Cortes-Reynosa et al, 2009). However, the detailed mechanism by which the internalized EGFR proteins are routed to the nucleus through the NPC is still unclear (Wanget al, 2010). To date it seems inappropriate to ascribe signaling importance only to second messenger pathways including mobilization of kinase and inducible transcription factor to explain nuclear effects of these factors (Rosenfeld and Hwa, 2009). These second messenger cascades, including JAK and STAT, are stimulated by so many ligands (Piwien-Pilipuk et al, 2002; Waters et al, 2006) that it is difficult to mediate the distinct cellular functions for each of these internalized molecules. Moreover, Lin et al (Lin et al, 1992) have previously reported that insulin is able to increase the mRNA levels of g33 and c-fos in cells treated with trypsin thus suggesting that the insulin-induced increase is mediated by a mechanism that is independent of insulin binding to its plasma membrane receptor and independent of the insulin receptor-associated kinase activity. As previously reported for insulin (Purrello et al, 1982; Soler et al, 1992), we used isolated nuclei in this study in order to study the effect of GH irrespective of its well-known effects mediated by the receptor pathway and the signal transduction pathway. As we previously demonstrated, GH is internalized via caveolae structures (Lobie et al, 1999), which means intracellular processes are likely to be direct effects of intact GH. The present approach provides the basis for additional studies to clarify the mechanism by which GH regulates gene expression. The variations in gene transcription were observed in the total absence of GH binding to the plasma membrane receptors and, therefore, generation of receptor-related signaling mechanisms.

In the present study, the kinetic of GH action on isolated nuclei reveals two effects: a direct stimulation of early gene by GH within 10 min of stimulation and later modifications of the same family as previously described for insulin by Lin et al (1992). In isolated nuclei, the consequences of GH effect are different with time, gene type and variation of expression.

If *in vivo* GH stimulated cell respiration, *in vitro* a direct, selective and dose-dependent effect of GH on isolated mitochondria was observed only on two enzyme (the inhibition of succinate dehydrogenase and cytochrome C oxidase activities) (Ardail et al, 2010). This latter inhibition was in contrast with the indirect, GH receptor-initiated stimulation of cytochrome C oxidase activity observed in GH-treated whole cells, showing that GH is specifically imported in mitochondria, where it operates a direct metabolic effect, independently of cell surface receptors and signal transduction.

Waters’ group reports (Conway-Campbell et al, 2007) that nuclear GHR correlates with high proliferative status as well as *in vivo* as *in vitro* studies, and that constitutive nuclear targeting of GHR causes a dysregulation of proliferative arrest and induction of cell cycle progression, resulting in oncogenesis. This appears to be a result of enhanced nuclear translocation of phospho-STAT5 in association with nuclear-targeted GHR. Other proliferative signals such as phosphatidyl inositol 3-kinase/Akt presumably contribute to the constitutive proliferation (Johnson et al, 2004). Since the complex IFN-γ receptor, visualized in the nucleus (Sadir et al, 2005; Lambert et al, 2000), can translocate phospho-STAT1 to the nucleus (Jeay et al, 2001), our data are in total accordance with this likely mechanism, although, we cannot exclude direct transcriptional actions by the nuclear receptor as reported for the epidermal growth factor receptor (Lin et al, 2001).

The period of nuclear accumulation of GHR after GH addition (Lobie et al, 1994; Mertani et al, 2003) and of GH (Mertani et al,1996) coincides with cell cycle initiation. In this way, in intact cells, associated to signal transduction, internalized GHR and GH increased gene expression in cellular growth and proliferation cluster. If similar effect was observed after internalization, this stimulation would be provided by another gene population. In isolated nuclei, where some GHR were present without GH addition, 30 min GH incubation revealed an accumulation of GH within nuclear matrix. This GH accumulation involves the modifications of the gene expression with time. However, gene expression seems decreased with time. Similar consequence has been observed in mitochondria as consequence of direct GH action on enzymes of respiratory chain (Sadir et al, 2000).

In conclusion, the effects of GH internalization on gene transcription, without signal transduction consequences, were characterized by a direct effect on gene expression, distinct from those stimulated by signal transduction. In similar functions, direct action of GH internalized in nucleus was observed on different genes compared to GH signal transduction from plasma membrane. Consequences of GH/GHR internalization in nucleus are in agreement with gene expression but correlation with signal transduction and internalization seems to be more complexe than two distinct pathways, or complementary effect on the same pathways.

